# Antlion larvae follow optimality rules in body orientation during sand-tossing

**DOI:** 10.64898/2025.12.02.691593

**Authors:** Francesca Lo Bello, Massimo De Agrò, Giorgio Vallortigara

## Abstract

Pit-building antlion larvae (*Myrmeleon* spp.) throw sand at prey that approach their pit using their head and mandibles as a backward-directed “biological catapult”. This mechanism restricts effective sand-tossing to a defined angular range behind the larva. When prey-like vibrations come from outside this range, antlions are known to reorient, but the consistency and rules behind this behaviour remain unclear. We tested whether larvae follow optimal turning strategies when reorienting toward a vibrating artificial stimulus placed in disadvantageous positions along the pit. Most larvae responded and often ended in an optimal sand-throwing position. Initial orientation strongly predicted turning direction: larvae rotated along the shortest angular path to reach the optimal sand-throwing orientation. Our results show that antlion larvae do not rely on rigid reflexes during prey capture but appear to integrate vibrational cues with spatial rules to minimize angular displacement. This reveals unexpected behavioural flexibility in a sit-and-wait predator with highly limited mobility.

## Introduction

Throughout evolution, predators have adopted diverse hunting strategies, including the widely used “sit-and-wait” approach (ambush strategy), in which they remain immobile until prey approaches. Among insects, antlions (*Myrmeleon* spp., order Neuroptera) represent a successful example of this predatory strategy (for review see (1,2)).

During their larval stage, antlion inhabit sandy or loose substrates in which they excavate an inverted conical pit. To do so, they move backward in progressively smaller circles (3), using their head and mandibles to dig the sand, ultimately ending at the centre of the pit (3–7). The pit’s structure makes it an effective trap because its sloped walls are close to the angle of repose (8), which depends on the grain size (9–12): steep enough to prevent prey from escaping, yet stable enough to avoid unwanted sand collapses (13,14). Once the pit is completed, the larvae position themselves at the bottom, buried beneath the sand with only the antennae and the anterior edges of the mandibles exposed (3,4,6,15). When a prey falls into the pit, the steep and unstable walls cause the prey to slide toward the centre where the antlion larvae, positioned under the sand, capture it by closing their jaws (1,4,6).

However, the prey may resist or delay its fall to the bottom of the pit. When this happens, antlions adopt a more active behavioural strategy (5): as soon as the prey is positioned roughly behind it, the predator buries its long, curved mandibles into the substrate to collect sand grains and throw them backward toward the prey (4,16), effectively using its head and mandibles as a “biological catapult” (1,4,5). This sand-throwing behaviour typically induces small sand slides that both distract the prey and reduce its grip on the pit’s surface, ultimately causing it to slip toward the centre, where it is captured by the antlion (13).

The antlion is remarkably precise in its sand-tossing behaviour. The larvae can in fact identify both the presence and crucially the position of an approaching prey thanks to the vibration it produces on the substrate (5). Thereafter, the antlion starts throwing sand directly towards the identified location, with an approximation of only ±5° (5).

However, this behaviour is mechanically constrained. Antlions can only toss sand toward the rear of their body, as their throwing mechanism is restricted to a backward arc. This range reflects the limited degrees of freedom of the head– prothorax joint, which makes it mechanically impossible for the larva to project sand toward the anterior sector (1).

These observations may suggest that the antlion acts as a preprogrammed insect-catching machine, engaging in automatic behaviours upon specific sensory triggers, with limited leeway for fine-tuning or adjustments. Devetak and colleagues (7) reported however that when antlions are facing frontally toward the preys, or after repeated failed sand tosses, they reorient their bodies, sometimes moving closer to probably gain a more advantageous orientation. However, this observation was only qualitative, so it remains unclear how common, flexible, or precise this behaviour is.

Here we designed an experiment to investigate the reorientation behaviour of antlions towards artificial, immobile prey, located in disadvantageous positions for sand-tossing. We hypothesize that antlions will consistently reorient their body upon prey detection, and that will do so following optimality rules: that is, following the minimal movement path to reach the first effective throwing position.

## Materials and methods

A total of 19 larvae of *Myrmeleon formicarius* (Linnaeus, 1767), commonly referred to as antlion, were collected from the wild in the Trentino region (North- East of Italy) during the end of summer 2024. Once in the lab, each animal was individually housed in a square wooden box measuring 8×8×4cm. Each box was filled for 3/4th of its size with dry sand with a grain diameter of approximately 1- 2 mm. Larvae were then left for at least 3 days, to allow them enough time to construct their pitfall traps. The boxes were placed in a room with access to natural light through a window, which was left open for the whole duration of the experiment to maintain the natural day/night cycle.

### Experimental procedure

After ensuring the antlion had properly constructed its cone, a video camera Flir GigE (resolution 1280×720, 30fps) was placed 40cm above the box. A LED lamp was placed above the pit to provide light for the recording. We constructed a simulated prey by taping three PVC bristles to a vibrator (Satisfyer Dual Love Connect App). The vibration intensity was set at the minimum level for which the contact of the PVC bristles with the sand was sufficient to elicit the antlion reaction (most of the times), but not strong enough to disrupt the sand pit.

At the beginning of each trial, the PVC bristles were put in contact with the sand at the edge of the pit, at a random location across the circumference (Fig. 1). The vibrator was then moved up and down such that the tip of the PVC bristles slid along the radius of the pit, ensuring the maintenance of a fixed angular position relative to the centre, for a total of 1 minute (see SI1). After this time, a single ant was dropped in the pit as food reward, to ensure the animal survival and maintain motivation. The animal was then left to rest for the day, and the experiment proceeded with the next animal. If during the stimulation the antlion did not produce any response, the video was excluded from the analysis. All trials were conducted between 10:00 and 17:00. Each antlion was tested a maximum of 5 times, with at least 24 hours between each trial.

**Fig. 1.**
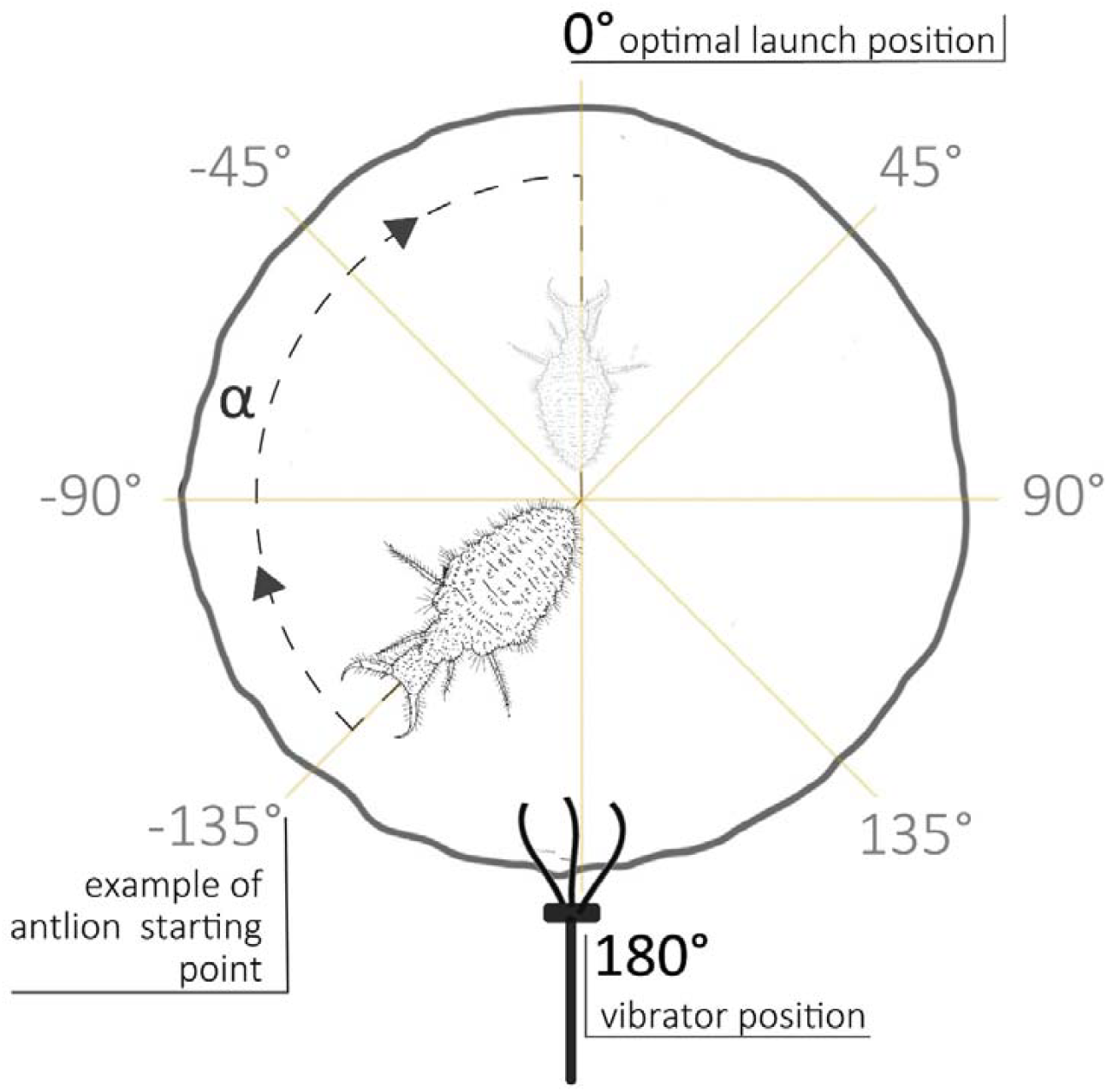
Schematic representation of the antlion pit showing the optimal sand-throwing position at 0°, the vibrator emitting vibrational stimuli placed at 180° and an example of the larva’s initial and final orientation. The angular displacement α represents the angle between the larva’s starting position and the optimal launch direction.

### Statistical analysis

A total of 137 trials were performed. The antlions produced a response in 45 of them, corresponding to 32,85%. From each video, we set the angular position of the PVC bristles point of contact relative to the antlion position as 180° (Fig. 1). Consequently, we recorded the initial orientation of the antlion by observing the very first throw upon stimulation. Orientations counterclockwise with respect to the 0° angle were set as negative values, while orientations clockwise rotated with respect to the 0° angle were set as positive values. These angular positions were approximated to the nearest 45°, for a total of 8 possible initial conditions (0°, 45°, 90°,135°,180°, -135°, -90°, -45°). We then followed the antlion behaviour and tracked its movement below the sand and its orientation. By the end of the trial, we recorded the final orientation (often being 0°, directly opposite to the PVC bristles position) of the antlion and which rotation direction it took to reach it (clockwise or counterclockwise).

Using the initial and final orientation of the animal along with the rotation direction, we calculated the total rotated angle α (again expressed as negative numbers for counterclockwise and positive numbers for clockwise rotations). The calculations were done in Python 3 (17), using the packages Pandas (18) and Numpy (19).

Statistical analysis was performed in R (20). We modelled the data using a generalized mixed effect linear model using the package GLMMtmb (21) using a gaussian error structure. The total rotated angle was set as the dependent variable, while the initial angle of the antlion was set as the predictor. Antlion identity was used as a random intercept. We then checked the effect of the predictor through an ANODA using the package car (22), and then performed a post-hoc, Bonferroni corrected analysis with the package *emmeans*(23). According to our hypothesis, antlions that started the trials correctly oriented with respect to the artificial prey position (at 0°, directly opposite to the prey) should not reorient, and so we would expect in this case a rotated angle of 0°. Antlions starting misoriented with a positive angle starting position should rotate counterclockwise in order to correctly orient themselves, and as such we should observe negative rotated angles. The reverse is true for initially misoriented antlions with a negative angle starting position. If instead the antlions rotate in a random direction until reaching optimality, the average rotated angle will be not statistically different from 0, having a mix of positive and negative rotation value. As a third possibility, antlions may show a lateralized preferential rotation direction. In that case we should observe either a positive or negative average rotated value, independent of the initial orientation.

## Results

Raw data and analysis are available in SI2. Of the 45 finally recorded trials, 12 had the antlion already perfectly oriented (0°) with respect to the artificial prey, while 5 were oriented oppositely (180°), and as such we had no expectation for any given rotation direction. Of the remaining 28, 12 were initially positioned at positive angles, and as such we expected a counterclockwise rotation, and vice- versa for the other 16. We excluded from the analysis the 5 antlions oriented at 180° and tested the effect of the initial orientation angle on the signed rotated angle. The ANODA revealed a significant effect of the predictor (χ^2^=5.326, p=0.021).

Of the 5 trials where animals that started at 180°, opposite to optimal orientation, 1 reoriented but we were unable to determine the direction due to it happening below the sand, 3 rotated clockwise and 1 rotated counterclockwise.

Specifically, the post-hoc analysis revealed a negative effect of the starting angle on the rotation amount (trend=-0.453, SE=0.196, t=-2.308, p=0.0269). This means that the more the starting angle is >0°, the more the animal tends to rotate counterclockwise (negative angles of rotation) to reach it, and vice versa.

Indeed, looking at the individuals’ behavior, only 1 out of 16 animals positioned at a negative angle rotated counterclockwise (while 5 did not rotate at all), and only 2 out of the 12 positioned at a positive angle rotated clockwise (while only 1 did not rotate; Fig. 2).

**Fig. 2.**
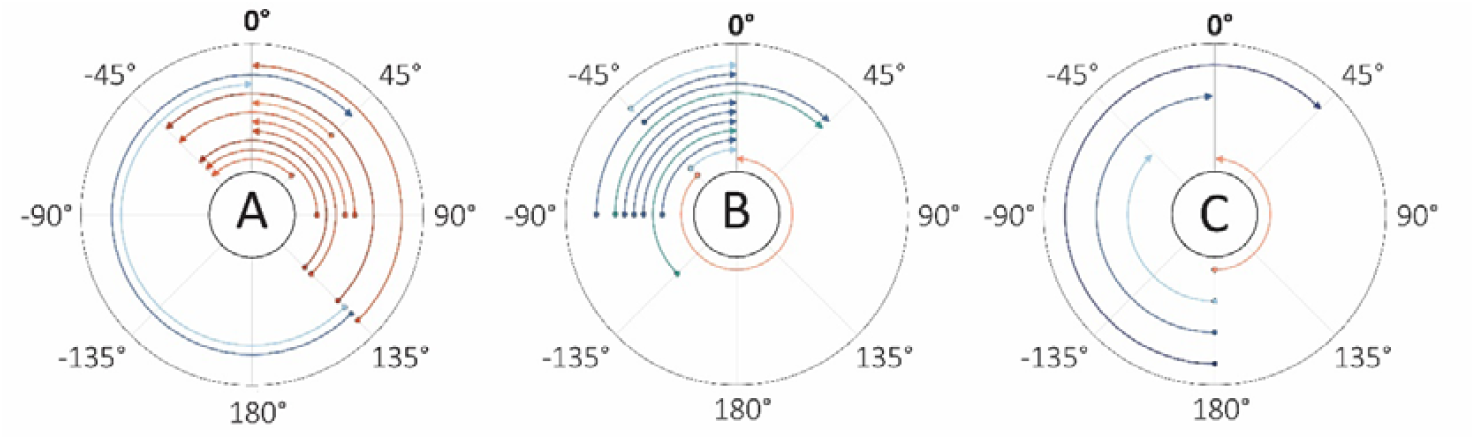
Each graph shows antlion rotation starting from a different angle: positive (A), negative (B) and 180° (C). Arrows indicate rotation direction and distance: red arrows correspond to counterclockwise rotations, blue arrows to clockwise rotations, with color intensity proportional to the angular displacement. 0° indicates the optimal launch position; 180° indicates the neutral position, from which the angular displacement to 0° is the same clockwise or counterclockwise.

Forty-one antlions ended the trial oriented in an advantageous throwing position, between +45° and -45°, equalling more than 90% of the trials (28 finished at 0°). In the 4 trials where subjects did not finish the trial in an optimal orientation, they did not ever reorient.

Of the 5 trials where animals that started at 180°, opposite to optimal orientation, 1 did not reorient, 3 rotated clockwise and 1 rotated counterclockwise. Even given this apparent imbalance, the sample is too small to suppose a rotational preference in the absence of an optimal choice.

## Discussion

Our results show that antlion larvae do not turn randomly or perform a purely automatic behaviour but follow a consistent, non-random turning strategy to reorient toward the prey-like vibrational stimuli.

Rather than rotating indiscriminately or following a fixed directional bias, larvae choose consistently the shortest rotational path to reach the optimal launch position.

Individuals starting on the negative side of the stimulus axis rotated mainly clockwise, whereas those starting on the positive side rotated predominantly counterclockwise, indicating that initial orientation strongly influences turning direction.

The observed relationship between initial orientation and rotation direction suggests that larvae integrate spatial information from vibrational cues with a rule of angular minimization. Furthermore, larvae positioned exactly at 180°, where the angular displacement required to rotate clockwise or counterclockwise is equal, showed a preference for counterclockwise rotation (the sample size of only 5 trials is too small to hypothesize a rotational preference in the absence of an optimal choice).

An interesting observation emerging from our experiment is that larvae often initiated sand tossing even before achieving an optimal throw-sand-position. We know that the trigger for initiating sand throwing is the detection of mechanical vibrations (1,13,24,25): antlions react immediately upon detecting prey-like cues but, if they find themselves misaligned with respect to the prey, they subsequently adjusts by rotating toward the appropriate orientation. We hypothesize that this failure might serve as the cue for initiating reorientation. Thus, the sequence may involve two distinct steps: an immediate vibrationally driven response, followed by a secondary, potentially error-driven correction when the first tosses prove ineffective. Although this remains speculative, it offers a plausible mechanistic framework for the observed behavioural pattern.

The results challenge the intuitive view of the antlion as a purely reflexive, pre- programmed ambush predator, revealing a previously unrecognized aspect of their predatory behaviour: the reorientation for sand-tossing shows flexibility and involves more than a simple stimulus-response chain. Our findings have broader implications for the study of decision-making in “sit-and-wait” predators. Optimal reorientation strategies could be a common feature among pit-building species and other ambush predators, allowing them to maximize efficiency while minimizing energy expenditure (26). Investigating such strategies across taxa could shed light on the evolution of spatial problem-solving and motor planning in animals with limited mobility.

## Supporting information

Supplemental Information 2 - Raw data and analysis

Supplemental Information 1 - Example Video

