## Supplemental Information 2 - Raw data and analysis for "Antlion larvae follow optimality rules in body orientation during sand-tossing": analysis.R

```
library(data.table) #To deal with big datasets
library(glmmTMB)
library(emmeans)
library(car)
library(DHARMa)
library(ggplot2)
library(parallel)

set.seed(12345)

## loading files

path <- 'C:/Users/massimo.deagro/Documents/Experiments/Antlions_FollowingPID/analysis/toPublish/antlion_degree.csv'


#### OPEN DATA ####

full_data <- fread(path)
full_data$trialn <- as.numeric(full_data$trialn)
head(full_data)

summary(as.factor(full_data$rotated_start_180))

sum(full_data$angular_distance>0)
sum(full_data$angular_distance<0)
sum(full_data$angular_distance==0)

full_data$starting_pos <- NaN
full_data$starting_pos[full_data$rotated_start_180>0] <- 'Positive'
full_data$starting_pos[full_data$rotated_start_180<0] <- 'Negative'
full_data$starting_pos[full_data$rotated_start_180==0] <- 'Aligned'
full_data$starting_pos[full_data$rotated_start_180==180] <- 'opposite'

sum(full_data$angular_distance>0)
sum(full_data$angular_distance<0)
sum(full_data$angular_distance==0)

table(full_data$ROTDIR, full_data$starting_pos)

table(full_data$rotated_end_180, full_data$rotated_start_180)

no180<-subset(full_data, full_data$rotated_start_180 != 180)
only180<-subset(full_data, full_data$rotated_start_180 == 180)


m0 <- glmmTMB(angular_distance ~ rotated_start_180 + (1|subj), data=no180, family = gaussian)

t.test(only180$angular_distance, mu = 0, alternative = "two.sided")

t.test(full_data$angular_distance, mu = 0, alternative = "two.sided")

Anova(m0)

et <- emtrends(m0, ~ 1, var = "rotated_start_180", type = "response")
test(et)

e <- emmeans(m0, ~ 1, type='response')
test(e)

ggplot(no180, aes(x=rotated_start_180, y=angular_distance))+
  geom_jitter(width = 5, height = 0)+
  geom_smooth(method = glm)
```
